# How competition for funding impacts scientific practice

**DOI:** 10.1101/2022.07.30.502158

**Authors:** Stephanie Meirmans

## Abstract

In the research integrity literature, funding enters in two different ways: as elevating questionable research practices due to perverse incentives, and as being a potential player to incentivize researchers to behave well. Other recent studies have emphasized the importance of the latter, asking funding experts. Here, I explored how the impact of competitive research funding on science is being perceived by active researchers. More specifically, I have conducted a series of group sessions with researchers in two different countries with a different degree of competition for funding, in three disciplinary fields (medical sciences, natural sciences and the humanities), and with researchers in two different career stages (permanent versus temporary employment). Researchers across all groups experienced that competition for funding shapes science, with many unintended questionable side effects. Intriguingly, these questionable effects had little to do with the type of questionable research practices (QRP’s) typically being presented in the research integrity literature. While the notion of QRP’s focuses on publications and assumes that there would essentially be a correct way to do the science, researchers worried about the shaping of science via funding. According to my session participants, rather than ending up as really being wrong, this shaping could result in predictable, fashionable, short-sighted, and overpromising science. And still, this was seen as highly problematic: scientists experienced that the ‘projectification’ of science makes it more and more difficult to do any science of real importance: plunging into the unknown or addressing big issues that would need a long-term horizon to mature.

## Introduction

There seems to be a crisis in science: surveys have recently found that many researchers perform questionable research practices (Bouter et al., 2016; Kaiser et al., 2021; Xie et al., 2021; Gopalakrishna et al., 2022). For example, they submit to selective reporting, p-hacking, and HARK-ing in order to score good publications (Bouter, 2020, following Wichters et al., 2016). Many research integrity scholars assume that it is the increasingly competitive nature of science, and in particular the need for high-impact publications and funding, which may be the main driver for individual researchers submitting to questionable research practices (Martinson et al., 2005, 2006; Bouter, 2020).

While for a long time there had been a focus on the individual researcher behaving badly, the focus in the research integrity debate has in recent years shifted away from individual responsibilities (and spectacular cases of fraud) to aspects of scientific communities and research climate. For example Zwart and ter Meulen (2019) have urged to investigate how universities and funders could help fostering research integrity.

Funding thus enters nowadays in two highly different ways into the discussion around research integrity: on the one hand, pressures on researchers to obtain funding are seen as potentially elevating questionable research practices. On the other hand, funders are also seen as potential agents to foster research integrity. How does this play out in practice, and what happens according to whom?

Labib and her colleagues (2021; see also Meijlgard et al., 2020) have recently made a first step to investigate how funding experts envision that they could help fostering research integrity. Labib et al. (2021) established eleven themes from the RI literature with regards to funding, and then asked funders about the significance of each theme. Using surveys, Labib and colleagues could identify which themes funders themselves would find most important to enhance responsible science. The top three themes that emerged in this way were “dealing with breaches of RI, conflicts of interest, and setting expectations on RPO’s (= research performing organizations)” (Labib et al., 2021). Funders were thus also seen as being able to impose requirements on research organizations with regards to research integrity of their employees (see also Roje et al., 2021 for a similar finding emerging from funding experts in focus group interviews).

What is currently lacking is the perspective from active researchers: how do researchers themselves experience the impact from funders on questionable research practices? And how does higher or lower competition factor into this? In this study, I investigated how active researchers experience the impact of competitive funding on their research, in a high versus in a low competitive setting.

## Methods

### Study design

Initially, I wanted to test the following hypothesis: does competitive research funding increase questionable research practices? This hypothesis was explored in an experimental design via doing group sessions with active researchers in two different countries: one country with a high degree of competition for research funding, and one country with a low degree of competition for research funding. The Netherlands was chosen as the ‘high-competition’ country (with grant success rates of 20-30%), and Switzerland as representative of a relatively ‘low-competition’ country (grant success rate 50-60%) (according to the Swiss Science Council and the Rathenau institute, personal communication). In addition, I also compared across different disciplines (natural sciences, medical sciences) and ‘seniorities’ in career stage – the idea being that juniors might be under higher pressure. Before performing the group sessions, I also conducted a couple of ‘pilot’ single interviews to gain a better understanding of what are the issues at hand and to help shape the group sessions.

### Insights from (pilot) single interviews

From the first couple of these pilot single interviews, mostly conducted in Switzerland in 2017, it quickly became clear that many researchers did not see a direct connection between competitive research funding and what are called ‘questionable research practices’. This did not mean, however, that my interviewees did not see any questionable effects of competitive research funding on doing good science. My interviewees told me many, and apparently for them quite serious, problems. However, these problems were often of a quite different nature than what is typically being captured under ‘questionable research practices’ in the research integrity literature.

In addition, it became clear that not all questionable research practices play a role in all disciplines. That this is indeed the case was also recently confirmed in a large-scale survey study, in which humanities scholars attested ‘not applicable’ to a large range of questionable research practices (Gopalakrishna et al., 2022). My interviewees also told me that it can even be the case that what is called a questionable research practice in one discipline can be a virtue in another (see also Ravn & Sørensen, 2021 for a similar finding): for example, diverging from an original research question is a virtue in the humanities but a vice in a medical study. There thus seemed to be a serious problem with the original research design of my study; it seemed to not allow gaining a universal understanding of the effects of competing for research funding on questionable/ responsible research practices.

Due to the insights gained during these pilot interviews, I decided to shift the original research question in the follow-up group sessions to a more open but rather simple question: “How does competitive research funding affect science (in good or bad ways)?” This question was accompanied by a follow-up question on what could be done better (results will follow in another publication).

### Participant selection and group session details

The sessions were conducted in 2017 in Switzerland and in 2018 in the Netherlands. In each country, six group session interviews were conducted. The groups consisted of typically 4 or 5 researchers, with a minimum of three in one case (natural science senior NL), and a maximum of seven researchers in another case (medical sciences senior CH). These researchers were grouped by scientific domains (natural sciences, medical sciences, and humanities) and career status. Career status was distinguished as ‘junior’ (=temporary employment) or ‘senior’ (=permanent employment)^1^. This made a total of twelve group sessions with in total 57 persons. Participants were recruited via personal networks as well as via Dutch and Swiss university websites and the website of the Royal Netherlands Academy of Arts and Sciences.

I checked session participants for their experience with funding ahead of the sessions and noticed that the recruitment strategy resulted in a high number of experienced researchers with funding. An overwhelming number of senior researchers had received multiple types of funding in the past (both via national funds, but many also had received EU funding, including an ERC for several participants). Many seniors had additional experience with participating in funding reviewing panels, at both the nationally and the international level, and including for ERC.

Each session took 3.5 hours. At the beginning of each session and via the invitation email researchers were familiarized with the background of the study (QRP’s) as well as with the idea that they could also explore other impacts of competition for funding on science, including in a positive sense. Written informed consent was obtained from all session participants. Researchers in the sessions then made extensive use of a digital tool called “Meetingsphere”. This tool is designed to allow anonymized digital interaction between session group members (https://www.meetingsphere.com). The tool was chosen due to the sensitive nature of the question, allowing honest answers regarding research integrity problems. At least half of the session time was spent on the following question: ‘How does competitive research funding affect science (in good or bad ways)?’ Group members were first allowed to type their answers into the digital system. After saturation of commenting (typically after 10-15 minutes), the system was opened for digital commenting (again around 10 minutes), followed by extensive oral discussion. One of the groups (Swiss natural science seniors) ended up with oral discussion only due to the delay of one of the session participants. My analysis for this paper focused on the digital session reports only, and the Swiss natural science group (with four participants) was therefore excluded from the specific analysis presented in this paper. My analysis here is thus based on the input of in total eleven group sessions and a total of 53 session participants.

### Analysis of session reports

I used a grounded thematic analysis in several rounds to analyze the Meetingsphere reports. I cross-checked identified themes in the first round with three other researchers, with analyses largely overlapping. In subsequent rounds, I refined the themes and split some into sub-themes.

## Results

Session participants were prolific in providing comments using the digital tool ‘Meetingsphere’, both with regards to initial own comments and in reaction to others’ comments. Via my thematic analysis, I identified a couple of main themes and subthemes that researchers addressed regarding the impact of competitive research funding on science in good or bad ways. Main themes consisted of: (1) The impact on how science is being shaped due to the competition for funding, (2) The impact of grant writing on research time, (3) The impact of publication pressure on detrimental research practices.

### 1. Shaping science

By far most comments received were within this category (262/ 317). These comments focused on how science is being shaped in practice via funding, and how this influence is being perceived and experienced. Importantly, these impacts are not seen as resulting in essentially wrong or sloppy science. Typically, the impact is experienced due to funder interventions, in both positive and negative ways. However, the negative often outweighs the positive. What typically happens is that researchers do understand and appreciate that funders select projects based on certain features, and that they intentionally shape funding calls and schemes in particular ways (positive). However, funder interventions can have unintended side-effects, and these can then be experienced as problematic by researchers (negative). Below, I provide an overview of the perceived impacts in subthemes. While some subthemes present positive and negative effects in a more balanced way, others show that the effects are predominantly experienced as negative. I also provide the number of comments within each subtheme, to give a sense of how much attention there was for each of the subthemes.

### Impact on science via peer review

(61 comments)

There were many comments on how funder peer review impacts science. Many researchers stated that competitive research funding should in theory increase overall quality in science by selecting the best proposals and scientists. Some Swiss researchers also thought this is indeed the case in practice.

*funding is brought to the best research ideas and best people* (nat jun, CH)

One Dutch researcher commented upon that success in funding acquisition often means future successes in gaining funding as well. This researcher was neutral about the effects on science via such a process: *I do not know whether it’s good or not*. (hum jun, NL).

There were a few comments on the positive effects of the competition on research practice. For example, one Swiss medical senior scientist said that *it improves research quality*. Researchers across countries and disciplines also expressed that projects that are submitted to funders typically have been thought through and tend to have solid methodologies. The feedback of reviewers can additionally help to improve the research, two Dutch natural senior scientists thought.

However, other – in particular Dutch - researchers perceived that while this is how it should work in theory the practice looks different. One important problem is that peer-review highly depends on the reviewers and the committee/ panel, and that these can be biased. Many told us that their comments were based on personal experiences, and negative experiences with biases in peer review led some Dutch senior researchers to state that peer review does not work anymore.

Humanities scholars (in both countries) thought that there is a deep problem because reviewers and panels can be biased if they represent certain research schools or fields. Such biases can even lead to a competition between scientific disciplines:

*how to avoid that competition between projects turns into competition between disciplines?* (hum sen, CH)

In the Netherlands, there was a specific problem with clustering of social sciences and humanities into one program. Due to disciplinary differences of what good research might mean several humanities scholars felt they had less chances to gain funding. One humanities researcher said that one would really need to address the question what science really is. The same type of bias was thought to play a role in gaining funding for medical qualitative research (where methods are different than in mainstream medical research).

*in the combined humanities & social-science boards, there is no understanding of what a humanities research project may look like*. (hum sen, NL)

On the positive side, one medical senior scientist expressed the view that an alternative system to the competitive research funding system might either not exist or be worse. In addition, several younger and older Swiss and Dutch humanities researchers mentioned that funding/ peer review can also enable to escape a limited home environment.

*Young researchers have the chance to free themselves from their home institutions by applying for funding and thus gain access to other cultures, ways of doing science*. (hum jun, CH)

### Impact on novel and risky science

(61 comments)

Researchers submitted many comments on this topic, across all 11 groups. The comments were predominantly negative. Many expressed that while funders often aim to fund innovative and risky projects, the opposite typically happens in practice. One Dutch researcher commented that the *‘rhetoric of innovation and breakthrough’* does not reflect how most funding is awarded in practice (hum jun, NL). The reason for this is that research projects are designed to be funded, not designed towards what would be considered ‘the best science’, new and original science.

*in principle, good effort to support the best science, but the measures of success are in favour of “productive” science, not necessarily creative science* (med sen, CH)

*the competitive system only works for ideas and methodologies that are well established, well known, not for ideas and methodologies that are new and really original* (hum sen, NL)

One reason for this is that funders put too much emphasis on the track record of the researcher, meaning that one dares not to stray away too far from own disciplinary grounds and instead “plays safe”. It “*encourages researchers to take small steps in the development of research ideas instead of taking a larger risk and trying something completely different”* (med jun, CH). It imposes a “*disciplinary straightjacket”* (hum sen, CH) to the individual researcher, it encourages researchers to “*remain within areas in which you have already proven yourself with publications”* (hum sen, CH).

“*Changing fields is discouraged in the current structure of competitive funding, a characteristic that is not supportive of interdisciplinarity and innovation*.*”* (med jun, CH)

For science, this means that research will progress only in “*incremental steps”* (med jun, NL), while this may not be the best research: “*it probably leads to conservative research”* (hum jun, NL). And this might in the end be counterproductive to what good science should be all about: taking risks, venturing into the unknown.

### Impact on science via funder research agenda

(40 comments)

Many researchers across countries experienced that funders steer what kinds of research can be done; this is on the one hand positive because money can strategically be put into solving important challenges:

*It enables society and politics to focus scientific research on key societal challenges and problems. In this sense, it contributes to societal problem-solving*. (nat jun, NL)

However, most researchers across countries experienced under agendas as being problematic because they might not foster the best science. Swiss scientists also commented that it would be disastrous if the funder agenda would bias against doing basic research:

*Negative/comment: It would be disastrous if competitive funding schemes would push research away from fundamental science* (nat jun, CH)

Indeed, many senior Dutch natural scientists experienced just that, even though some also saw positive aspects in more applied ways of doing science.

*Negative: nearly all 100% fundamental project funding possibilities in NL are being eliminated. Even the Science Agenda is now funded with contributions from industry*. (nat sen, NL)

The same concerns held true for other types of valorization in the Netherlands: valorisation can take time away from doing core research work. Funder bias can also mean that bigger research fields or those with a higher applicability are more likely funded, which both natural and medical scientists across countries experienced as problematic.

### Impact on science via incentivizing collaborations

(29 comments)

Researchers frequently reported that funding has effects on collaborations. It typically fosters to collaborate, and many researchers regarded this in principle as positive. For example, one Dutch medical senior perceived this as good because *it helps to establish interactions and networks beyond the finally funded projects*. Another medical Swiss senior expressed that *the process of writing applications already has major impact on creating innovative idea and collaborations*.

However, many researchers also experienced that those collaborations often do not work well in research practice. This can be due to a variety of reasons, such as too large consortia, inter-disciplinary problems or feeling forced to collaborate. This can be problematic to a degree that collaborations have negative effects in practice. Often medical seniors uttered such skepticism about large consortia/ interdisciplinary multicenter collaborations. They in practice do not work well, they said, there are communications problems between disciplines, and they would *need better support and guidance* (med sen, CH). They can be forced upon you, and lead to a lot of *formal interaction without actual benefits* (nat jun, CH). In terms of certain collaborations, you could better do without in practice.

*forming strong consortia to increase chances; this can also be a disadvantage if you feel obliged to cooperate with groups for increasing chances on funding, but that will either just complicate the research process / feasibility or even be a disadvantage* (med sen, NL)

Such sobering practices can lead to dishonesty about collaborations in applications. One senior Swiss humanities researcher wrote that they often exist only on paper, are fake. Funding can also lead to confusing effects with regards to collaborations and team science, for example in the humanities which does not have a tradition of ‘team research’.

### Impact on science via research plannning

(20 comments)

Many researchers across groups expressed that applying for funding has a positive effect on thinking through, planning and structuring research. This can make researchers *think about next steps in your research* (med sen, NL) and *think carefully about* what to do and how to do it. Ultimately, this *might help to make [the research] more effective and more fruitful* (both hum jun, NL).

Some Dutch natural scientists also expressed that the need to apply for funding could even help to come up with new ideas and trigger new collaborations, for example with other groups with better skills. It can in practice also enable the researcher to spend time on thinking and getting up to date with the literature. However, Dutch medical senior researchers also perceived that the way good science should be done often is at odds with the way funding works:

*It also limits flexibility to change the design when needed or address additional question which appear more interesting on the way*. (med sen, NL)

One Dutch natural science researcher thought this is not so much of a problem in practice, because “*surely no one does exactly what is in the grant, right? You write a cool proposal and decide later what’s actually possible”* (nat jun, NL). Other researchers did feel forced to become dishonest in their grant-writing in order to circumvent this epistemic problem: *Bad: Science is per definition not predictable. Competitive funding forces you to predict your science, i*.*e. first do experiments than write the grant. Afterwards claim success because all your ‘predictions’ turned out to be true. This is often termed ‘pilot’-data* (med sen, NL)

*‘You have to have 2/3 of the paper already written to get the grant for the project’* (med sen, NL)

### Impact on research via length of funding period

(18 comments)

Another effect of funding on scientific practices was that grants typically are for shorter periods only – typically a couple of years. Such limitations can restrict the design of a project and lead to a focus on *short term deliverables* (med jun, NL). One Dutch senior natural science researcher experienced this effect as positive, and even thought that having such short-term funding could benefit long-term research lines in the end because the expectation of the release of data and new results stimulates you to work harder.

However, most researchers, across countries and disciplines, saw the impact of time-limited funding schemes as a potential danger for doing good science. They expressed that *it can be difficult to continue a line of research* (nat sen, NL) and that *long term research is being prevented (*nat sen, NL). The latter is a problem because *big societal problems require long term data* (nat sen, NL). It was obvious that many researchers considered research done over a long time as highly valuable but endangered by funding practices. In the humanities, some scholars feared short periods of time would not even allow to do any significant research at all. One compared short-term research in the humanities with building *pre-fab houses, but no cathedrals* (hum sen, NL)

*Most competitive research funding is project based and 3-4-5 years duration. It is highly questionable whether this system adequately supports academic research in the humanities since this research often takes much longer period of times to mature*. (hum jun, NL)

Several senior researchers also reported short-term funding as leading to hectic research due to the time pressure, sometimes even leaving some of the gathered data to be un-analysed in the end.

### Impact on science via strategic grant applications

(18 comments)

Many researchers across countries, seniorities and disciplines mentioned that researchers strategically tailor their research ideas, topics, design and methods to what they think will likely receive funding. This can mean submitting to funder ideas and programs at the expense of own interest and ideas, which can imply impoverishment of science:

*Research projects are designed to be funded what might be different to research projects with very innovative and “unusual” ideas* (med jun, CH)

It can also mean tailoring research to fit into funder requirements and previously successful templates or restricting design of a project to the specific guidelines set out by the funder. One researcher puts very clearly that *The first question a researcher will always ask him/herself when writing a grant proposal is: “What is the right strategy to get the grant?”* (nat jun, CH). Such fitting of research to funding requirements can then be followed up in reality (or not). As a result, one researcher feared decreasing diversity in science:

*It makes everyone jump through the same hoops, everyone has to meet roughly the same criteria. In this sense it works against diversity in the Dutch science system*. (nat jun, NL)

It can also mean strategically generating income, part of which will be used to fund the ‘real’ research of interest:

*sometimes large research proposals may be written to generate income, only a small fraction of which (the spoils) are used to fund basic research that the principal investigators are actually interested in* (med jun, NL)

### Impact on science by feeling the need to write a ‘sexy’ proposal

(15 comments)

Across countries, seniorities and disciplines, researchers experienced that supposedly sexy, fashionable, topics and research proposals are more likely to be funded:

*Funding calls for ‘sexy projects’* (med jun, CH)

However, researchers did not think that these kinds of projects are typically of high scientific value because it does not focus on good science. And though it can also have positive effects of building trends, it can also have the problematic side effect to reduce diversity in scientific topics, disciplines, methods:

*skew/select specific trends, and then everyone jumps on the bandwagon - positive effect is that this can rapidly accelerate a promising direction, negative effect is that it creates bubbles/echo chambers which suck funding away from other directions (since the ultimate pool of money is not infinitely increasing*. (nat jun, CH)

### 2. Impact of grant-writing on research time

(28 comments)

Another theme expressed by junior and senior Dutch, and junior Swiss, researchers in all fields was that the constant need to apply (or act as reviewer) for funding is extremely time-intensive and distracts from time spend on and care for ongoing research. For junior researchers, this can mean spending a considerable amount of time during a given project on writing an application for the next one. This means that one cannot invest sufficiently in the project one is currently undertaking. And senior researchers, many comments claimed, often do *more* grant-writing or grant-evaluating than research. This problem is particularly severe if funding rates are low:

*Takes up a lot of time and effort that basically goes to waste if the project is not funded - problem especially when, as is the case with NWO, the chances of getting funding are so low. (bad thing)* (hum sen, NL)

Some Dutch junior humanities scholars actually doubted the overall value of such a funding system - due to the time investments that currently need to be made. Also the associated administration costs are thought to be too time intensive by some Swiss researchers, and they said that this time could better be used to do research.

There were only a handful positive effects of funding on time management being reported. These comments were exclusively brought forward by senior natural science and senior medical researchers across countries. One Swiss medical researcher for example thought that c*ompetitiveness can trigger[s] an environment that stimulates the investment of effort (time, thought, hard work)*. One Dutch senior natural scientist thought that the need to devote some time towards writing grants can provide you with time to do some creative thinking.

### 3. The impact of publication pressure on detrimental research practices

There were comparatively few comments provided within this theme, a mere 9% of all comments. Below, I distinguish between comments mentioning questionable research practices and sloppy science (5%), and those stating scientific malpractices (3,5%).

#### Impact on questionable research practices and sloppy science

(16 comments)

There were some remarks on the occurrence of questionable research practices. Interestingly, statements regarding negative effects through publication pressure were made mostly by junior Swiss researchers in the natural sciences and the humanities, though there was one statement by a young Dutch humanities scholar as well. This finding stands in contrast to the hypothesis that researchers in a country with a higher funding rate (and thus supposedly less competition) should put a more relaxed focus on publications.

Junior researchers expressed for example the view that the following questionable publication practices are taking place due to the publication pressure: *splitting research into minimal publishable pieces, self-plagiarism, hasty and not fully careful analyses, etc*. (hum jun, NL). Another researcher thinks that *junior researchers may be tempted to write papers with controversial views* (hum jun, CH), or submit to *exaggerating impact both in proposal and in publications (overhyping)* (nat jun, CH).

Several Swiss natural science and humanities juniors emphasized that publication pressure could result in haste versus care. Interestingly, junior researchers then assumed that this is predominately problematic for reviewers who might need to put a lot of effort and time into correcting this. At least some researchers thus apparently thought that sloppy research would eventually get corrected via journal peer reviewing.

One researcher mentioned that publication pressures are not primarily exerted by the funding system but rather by the academic career system:

*In my opinion this [rapid publication versus careful analysis] is a problem related to extreme weight given to publication record when academics apply for positions*. (nat hun, CH)

On the other hand, several – mostly senior – medical and natural sciences researchers across countries expressed that the publication pressure which the system exerts can also be positive because it ensures that papers are eventually being published.

#### Impact on research misconduct

(11 comments)

Only four of the in total 53 interviewees commented that competition for funding could result in research misconduct, three of which were either Swiss or Dutch medical senior scientists. One of the Swiss ones for example said that *the high pressure for success obviously fosters the danger of data fabrication, which is extremely difficult to control* (med sen, CH). The reasons for fringe behaviour, another Swiss said, may be extreme competition amongst PI’s.

However, the Dutch medical scientist commented that if bad practices indeed occur, the problem may have to be viewed in a much broader perspective than funding per se. One would need to consider also “*researcher’s careers, positions, salaries etc”*. Because these aspects are based on the same criteria. Interestingly, the same scientist also admitted that occurrence of bad practices in his/her case were mainly based on hearsay and not on own experiences. They were thus essentially speculations. It is then interesting to note that the fourth person mentioning a potential occurrence of severe research misconduct formulated the comment as a question:

*if your livelihood depends on it, doesn’t it seem very understandable to tweak the results of your study so to increase the chance of that high impact paper that will help you get your next funding?? (nat sci jun, NL)*

In this group, the four other junior Dutch natural scientists all individually reacted to such an (in their eyes) extreme view of unethical behaviour, even though they admitted that scientists may behave in strategic ways and thus do things too sloppy or somewhat biased.

*I think “cheaters’” is maybe a bit too strong. I would say that the funding system stimulates “strategic behaviour”, i*.*e. behaviour to maximize the quantifiable output of research. (nat sci jun, NL)*

## Discussion

Researchers involved in my study experienced that competition for funding has a drastic effect on scientific practice. While some of these effects are positive, most effects are perceived as problematic. Those problematic effects, however, were of a quite different nature than what typically is perceived as questionable research practices (QRP’s) in the research integrity literature. According to session participants, competitive research funding did not have a big impact on detrimental scientific practices (a mere 9% of the comments provided). Publication pressure was experienced more as a general phenomenon in academia. Contrary to expectations, it was junior researchers in the *low*-competition country which at all connected funding with publication pressures.

The effects on science which researchers perceived as most important (91% of comments) were direct effects on science introduced by funding. Most of these effects were expressed by all session groups. Such effects were typically of a much broader nature than performing single publications or studies in a correct manner. The underlying mechanism seems to be the following one: funders aim to incentivize researchers to do good science. For example, by asking for explicit proposals they should select and provide money for the best science. Researchers are also being pushed to valorise, to broaden their perspective by collaborating in bigger teams, or to show that their projects are feasible. And while researchers do appreciate and value these intentions, they often feel that they have questionable unintended side effects in practice. Selection via peer review can have questionable effects of decreasing diversity. Feasibility often results in non-risky predictive research. Valorisation bends away from putting sufficient care into the core research. Working in teams can turn out to be extremely difficult and diminish individual researcher maturation. I would suggest that many of such intended and unintended aspects fell under the umbrella of the ‘projectification’ of science induced by funding (see also Felt, 2021a). Via shaping science into ‘projects’, funding has unintended side effects of at least some science to become predictable, boring, short-sighted, fashionable and/or overpromising. Researchers wooried that this might make it difficult to do good science that really matters: plunging into the unknown or addressing big issues that would need a long-term horizon to mature.

High competition for funding in the Netherlands seems to have exacerbated such unintended effects of funding in an interaction effect (but not QRP’s). The Netherlands does not only have a more competitive funding system, but is also steered by science policy to a much higher degree than Switzerland, with researchers experiencing less autonomy (Lepori et al., 2007), for example also with regards to valorisation (de Jong et al., 2016). This effect was visible in my findings: Dutch researchers were more vocal and experienced with negative side effects of strong science policies, such as little budget for basic science or aspects of valorisation. And I speculate that those effects overshadowed any effects of publication pressure with regards to the Netherlands (which is why I might have found a higher perception of publication pressure amongst Swiss junior scientists than Dutch ones). Swiss scientists seemed in comparison much happier with their funding system, and this went beyond pure aspects of lower competition (higher autonomy).

When looking at the scholarly literature beyond research integrity, none of my above findings on how funding shapes science is very novel or surprising. Scientists have over the years repeatedly pointed out that competing for funding impacts science in for them often worrying ways (starting as early as in the 1970s, see e.g. Brooks, 1978). There are a whole host of science policy and other studies addressing and discussing the relationship between details of competitive research funding and scientific practice. Topics include for example funder peer review and its biases (Bornmann & Daniel, 2006; Langfeldt, 2006; van den Besselaar & Leydesdorff, 2009), valorisation (Wallace & Rafols, 2015; de Jong et al., 2016), and risky versus conservative science (Guthrie et al., 2019; Veugelers et al., 2019; Ayoubi et al., 2021). More recently, some studies have started making a connection between this literature and the research integrity literature (Conix et al., 2021; Recio-Saucedo et al., 2022).

My study is novel in exploring the effects of competitive research funding bottom-up, showing that the current focus on QRP’s might misrepresent where actual problems with doing good science in connection with funding lie. Other studies of a comparable ethnographic kind have made similar findings with regards to what it would mean to do good science and what currently restricts it (Jerak-Zuiderent et al., 2021), also with regards to the impact of time and projectification (Felt 2021 a,b). But are the insights generated by my study still about research integrity per se? Hasn’t it in the end become, as above studies seem to suggest, more about science policy? Shouldn’t we rather strive for a more explicit demarcation of what research integrity actually is (Helgesson & Bülow, 2021)? However, other research integrity researchers also already emphasize that there needs to be a shift in focus from individual researcher responsibilities to aspects of the ‘system’ (Bonn & Pinxten, 2019; Bruton et al., 2020; Sørensen et al., 2021). In addition, what is currently understood under research integrity seems to depend already on whom you ask (Davies, 2019; Davies & Lindvig, 2021).

I would suggest that our goal in connection with funding should be to find out what the real problems on doing good and valuable science are – and ultimately, what issues funders and other science policy makers should address to improve the situation. Looking at this from several perspectives is certainly valuable. My findings are very different from the ones reached by Labib et al. (2021) and Roje et al. (2021), and also my recommendations would be different: Funders should reflexively re-evaluate some of the specifics of their funding schemes. And shouldn’t our main concern be about how we enable researchers to do good science?

## Acknowledgements

I thank all session participants for their time and for sharing their insights with me. I would also like to thank the Swiss Academies of Arts and Sciences, and in particular Roger Pfister, for hosting the sessions in Switzerland. I am also indebted to Gerd Folkers, whose guidance and contacts to Swiss scientists was of uttermost importance for this study. I also thank the Royal Netherlands Academy of Arts and Sciences, and Jean Philippe de Jong in particular, for helping to contact interviewees, hosting the discussion sessions in The Netherlands and engaging the Swiss Academies in the project. Furthermore, I would like to thank Herman Paul, Jeannette Pols, Barend van der Meulen and Peter van Hoesel for good advice throughout the project, as well as Danny van den Boom with helping to make the Meetingsphere sessions a success. The study was funded by a ZonMw grant, # 445001004.

## Footnotes

1 In the Results, the following abbreviations are being used: med = researcher in a medical field; nat = researcher in a natural science field; hum = researcher in the humanities; jun = junior; sen = senior; NL = researcher currently based in the Netherlands; CH = researcher currently based in Switzerland

## References

Ayoubi, C., Pezzoni, M., & Visentin, F. (2021). Does it pay to do novel science? The selectivity patterns in science funding. Science and Public Policy, 48(5), 635–648. https://doi.org/10.1093/scipol/scab031

Bonn, N. A. & Pinxten, W. (2019). A decade of empirical research on research integrity: what have we (not) looked at? Journal of Empirical Research on Human Research Ethics, 14(4), 338–352. https://doi.org/10.1177/1556264619858534

Bouter, L. (2020). What Research Institutions Can Do to Foster Research Integrity. Science and Engineering Ethics, 26, 2363–2369. https://doi.org/10.1007/s11948-020-00178-5

Bouter, L. M., Tijdink, J., Axelsen, N., Martinson, B. C. & ter Riet, G. (2016). Ranking major and minor research misbehaviors: results from a survey among participants of four World Conferences on Research Integrity. Research Integrity and Peer Review, 1, 17. https://doi.org/10.1186/s41073-016-0024-5

Bornmann, L., & Daniel, H.-D. (2006). Potential sources of bias in research fellowship assessments: effects of university prestige and field of study. Research Evaluation, 15(3), 209–219. https://doi.org/10.3152/147154406781775850

Brooks, H. (1978). The Problem of Research Priorities. Daedalus, 107(2), 171–190. http://www.jstor.org/stable/20024552

Bruton, S. V., Medlin, M., Brown, M., & Sacco, D. F. (2020). Personal motivations and systemic incentives: Scientists on questionable research practices. Science and Engineering Ethics, 26, 1531–1547. https://doi.org/10.1007/s11948-020-00182-9

Conix, S., de Block, A., & Vaesen, K. (2021). Grant writing and grant peer review as questionable research practices. F1000Research, 10:1126. https://doi.org/10.12688/f1000research.73893.2

Davies, S. R. (2019). An ethics of the system: Talking to scientists about research integrity. Science and Engineering Ethics, 25(4), 1235–1253. https://doi.org/10.1007/s11948-018-0064-y

Davies, S. R., & Lindvig, K. (2021). Assembling research integrity: negotiating a policy object in scientific governance. Critical Policy Studies, 15(4), 444–461. https://doi.org/10.1080/19460171.2021.1879660

de Jong, S. P. L., Smit, J., & van Drooge, L. (2016). Scientists’ response to societal impact policies: A policy paradox? Science and Public Policy, 43(1), 102–114. https://doi.org/10.1093/scipol/scv023

Felt, U. (2021a). Making and taking time: Work, funding and assessment infrastructures in inter- and trans-disciplinary research. In B. Vienni Baptista, & J. Thompson Klein (Eds.), Dynamics of inter- and trans-disciplinarity within institutions: Cultures and communities, spaces, and timeframes.

Felt, U. (2021b). In conclusion: The temporal fabric of academic lives: Of weaving, repairing, and resisting. In F. Vostal (Ed.) Inquiring into Academic Timescapes (pp. 267–280). Emerald Publishing Limited, Bingley. https://doi.org/10.1108/978-1-78973-911-420211022

Gopalakrishna, G., ter Riet, G., Vink, G., Stoop, I., Wicherts, J. M., & Bouter, L. M. (2022). Prevalence of questionable research practices, research misconduct and their potential explanatory factors: A survey among academic researchers in The Netherlands. PLoS ONE, 17(2), e0263023. https://doi.org/10.1371/journal.pone.0263023

Guthrie, S., Rodriguez Rincon, D., McInroy, G. et al. (2019). Measuring bias, burden and conservatism in research funding processes [version 1; peer review: 1 approved, 1 approved with reservations] F1000Research, 8:851. https://doi.org/10.12688/f1000research.19156.1

Helgesson, G., & Bülow, W. (2021). Research integrity and hidden value conflicts. Journal of Academic Ethics. https://doi.org/10.1007/s10805-021-09442-0

Jerak-Zuiderent, S., Brenninkmeijer, J., M’charek, A., & Pols, J. (2021). Good science: A view from within. Amsterdam: Amsterdam UMC, report, 36 p.

Kaiser, M., Drivdal, L., Hjellbrekke, J. et al. (2022). Questionable research practices and misconduct among Norwegian researchers. Science and Engineering Ethics, 28, 2. https://doi.org/10.1007/s11948-021-00351-4

Labib, K., Roje, R., Bouters, L., Widdershoven, G., Evans, N., Marušić, A., Mokkink, L., & Tijdink, J. (2021). Important topics for fostering research integrity by research performing and research funding organizations: A Delphi consensus study. Science and Engineering Ethics, 27, 47. https://doi.org/10.1007/s11948-021-00322-9

Langfeldt, L. (2006). The policy challenges of peer review: managing bias, conflict of interests and interdisciplinary assessments. Research evaluation, 15(1), 31–41. https://doi.org/10.3152/147154406781776039

Lepori, B., van den Besselaar, P., Dinges, M., Potí, B., Reale, E., Slipersæter, S., Thèves, J., & van der Meulen, B. (2007). Comparing the evolution of national research policies: what patterns of change? Science and Public Policy, 34(6), 372–388. https://doi.org/10.3152/030234207X234578

Martinson, B. C., Anderson, M. S., & de Vries, R. (2005). Scientists behaving badly. Nature, 435(7043), 737–8. https://doi.org/10.1038/435737a

Martinson, B. C., Anderson, M. S., Crain, A. L., & de Vries, R. (2006). Scientists’ perceptions of organizational justice and self-reported misbehaviors. Journal of Empirical Research on Human Research Ethics, 1, 51–66. https://doi.org/10.1525/jer.2006.1.1.51

Mejlgaard, N., Bouter, L. M., Gaskell, G., Kavouras, P., Allum, N., Bendtsen, A. K., et al. (2020). Research integrity: nine ways to move from talk to walk. Nature, 586, 358–60. https://doi.org/10.1038/d41586-020-02847-8

Ravn, T., & Sørensen, M. P. (2021). Exploring the gray area: Similarities and differences in questionable research practices (QRPs) across main areas of research. Science and Engineering Ethics, 27, 40. https://doi.org/10.1007/s11948-021-00310-z

Recio-Saucedo, A., Crane, K., Meadmore, K., Fackrell, K., Church, H., Fraser, S., & Blatch-Jones, A. (2022). What works for peer review and decision-making in research funding: a realist synthesis. Research Integrity and Peer Review 7, 2. https://doi.org/10.1186/s41073-022-00120-2

Roje, R., Tomić, V., Buljan, I. & Marušić, A. (2021), Development and implementation of research integrity guidance documents: Explorative interviews with research integrity experts. Accountability in Research. https://doi.org/10.1080/08989621.2021.1989676

Sørensen, M.P., Ravn, T., Marušić, A., Reyes Elizondo, A., Kavouras, P., Tijdink, J. K., & Bendtsen, A.-K. (2021). Strengthening research integrity: which topic areas should organisations focus on?. Humanities and Social Sciences Communications, 8, 198. https://doi.org/10.1057/s41599-021-00874-y

van den Besselaar, P., & Leydesdorff, L. (2009). Past performance, peer review and project selection: a case study in the social and behavioral sciences. Research Evaluation, 18(4), 273–288. https://doi.org/10.3152/095820209X475360

Veugelers, R., Wang, J., & Stephan, P. (2019). Do funding agencies select and enable risky research: Evidence from ERC using novelty as a proxy of risk taking. Extended Abstract submitted for the NBER SI SSF Workshop, July 2019.

Wallace, M. L., & Rafols, I. (2015). Research portfolios in science policy: Moving from financial returns to societal benefits. SPRU Working Paper Series (SWPS), 2015-10: 1–30. ISSN 2057-6668. Available at http://www.sussex.ac.uk/spru/swps2015-10

Wicherts, J. M., Veldkamp, C. L. S., Augusteijn, H. E. M., Bakker, M., van Aert, R. C. M., & van Assen, M. A. L. M. (2016). Degrees of freedom in planning, running, analyzing, and reporting psychological studies: A checklist to avoid p-hacking. Frontiers of Psychology, 7, 1832. https://doi.org/10.3389/fpsyg.2016.01832

Xie, Y., Wang, K. & Kong, Y. (2021). Prevalence of research misconduct and questionable research practices: A systematic review and meta-Analysis. Science and Engineering Ethics 27, 41. https://doi.org/10.1007/s11948-021-00314-9

Zwart, H., & ter Meulen, R. (2019). Addressing research integrity challenges: from penalising individual perpetrators to fostering research ecosystem quality care. Life Sciences, Society and Policy, 15, 5. https://doi.org/10.1186/s40504-019-0093-6

